# A DNA Sequence Imaging Approach to Predict Splice Sites Using Deep Learning

**DOI:** 10.1101/2025.06.24.661415

**Authors:** Espoir Kabanga, Seonil Jee, Arnout Van Messem, Wesley De Neve

## Abstract

We present a novel approach to splice site prediction using image-based deep learning, comparing the established Frequency Chaos Game Representation (FCGR) with our proposed Dinucleotide Fixed Color Pattern (DFCP) technique. Applied to donor and acceptor splice site sequences from *Arabidopsis thaliana* and *Homo sapiens*, DFCP consistently outperforms FCGR in accuracy, precision, recall, and F1-score when using a ResNet50 model. Visualization techniques such as saliency maps and Grad-CAM further demonstrate that DFCP produces more localized and biologically interpretable activation patterns. These findings highlight the critical role of sequence visualization strategies in enhancing deep learning performance and interpretability in genomic analysis.

## 1 Introduction

Over the past decades, researchers have explored a wide range of strategies to better understand and analyze patterns in DNA sequences. Among these, signal processing and numerical transformations have played a substantial role in shifting the view on DNA from a linear string of letters to a structured signal. For example, the work done in [1] introduced a large-scale feature analysis of genomic signals, while the work in [2] developed techniques to convert nucleotide sequences into complex-valued signals, allowing analyses in the frequency domain and revealing periodicities in coding regions and long-range correlations across extended genomic regions.

Building on this foundation, more recent approaches in graphical bioinformatics [3] have focused on transforming DNA sequences into two-dimensional (2D) visual representations. This shift has been driven in part by the success of deep learning [4], particularly convolutional neural networks [5, 6], in analyzing visual data. One of the earliest and most widely adopted methods for sequence visualization is Chaos Game Representation [7, 8, 9], which maps each nucleotide to a position in a fractal-like square to produce an interpretable visual signature of a DNA sequence.

An extension of this concept, Frequency Chaos Game Representation (FCGR), encodes the frequency of *k*-mers into a 2D matrix, resulting in an image where spatial patterns reflect compositional properties of the sequence [10, 11]. FCGR has proven useful for tasks such as classification and prediction, particularly when used in combination with deep learning [12, 13].

In this study, we propose a new visual encoding method, Dinucleotide Fixed Color Pattern (DFCP), which captures pairwise relationships between all positions in a DNA sequence. Unlike FCGR, which is frequency-based, DFCP creates a square image where each pixel represents a dinucleotide combination between two positions, mapped to a unique RGB color. This approach preserves both local and long-range dependencies in a consistent visual pattern.

To evaluate the effectiveness of the FCGR and DFCP visualization strategies, we applied them to the task of splice site prediction. Splice sites mark exon–intron boundaries and play a key role in gene expression [14]. Accurate prediction of splice sites remains a core challenge in computational genomics, as both local motifs and broader sequence context influence splicing decisions [15].

We used a ResNet50 model [16] trained on FCGR and DFCP images derived from *Arabidopsis thaliana* and *Homo sapiens* sequences containing annotated splice sites to evaluate how different visual encoding strategies influence predictive performance and model interpretability in the context of genomic sequence classification.

## 2 Materials and Methods

### 2.1 Dataset

We used donor and acceptor splice site sequences from *Arabidopsis thaliana* and *Homo sapiens*, obtained from the DRANetSplicer dataset [17]. Each sequence consisted of 402 nucleotides centered around a true or potential splice site. The dataset included both positive examples (true splice sites) and negative examples (non-splice sites).

For each category (*Arabidopsis thalina* donor, *Arabidopsis thaliana* acceptor, *Homo sapiens* donor, and *Homo sapiens* acceptor), we selected 35,000 positive and 35,000 negative sequences. These were converted into image representations using both the FCGR and the DFCP visualization techniques, resulting in 70,000 images per category for each visualization method.

We partitioned the data in a training set consisting of 50,000 images (25,000 positive and 25,000 negative), a validation set of 10,000 images (5,000 positive and 5,000 negative), and a testing set of 10,000 images (5,000 positive and 5,000 negative). The same partitioning was applied to both the FCGR and DFCP images to allow consistent and fair comparisons between the two encoding strategies.

### 2.2 Frequency Chaos Game Representation

FCGR builds upon CGR, a technique originally used to visualize DNA sequences as fractal-like patterns. In CGR, each nucleotide incrementally moves a point within a square by halving the distance toward a fixed corner assigned to that nucleotide, producing a spatial pattern that reflects the sequence composition and order. FCGR extends this idea by counting the frequency of all possible *k*-mers and mapping them onto a two-dimensional grid.

Each *k*-mer is assigned a unique position in a 2^*k*^ × 2^*k*^ matrix, where *k* is the *k*-mer length. Nucleotides are first converted into binary codes: A = 00, C = 01, G = 10, T = 11. For a given *k*-mer *s* = *s*_1_*s*_2_ *… s_k_*, its matrix coordinates (*x, y*) are computed as:

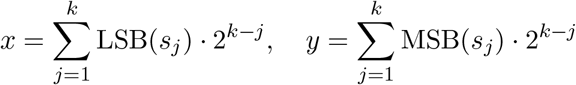

where LSB(*s*_*j*_) and MSB(*s*_*j*_) are the least and most significant bits of the binary code of *s*_*j*_.

We used *k* = 6, resulting in 64 × 64 matrices (4^6^ = 4096 possible *k*-mers). This choice provided a balance between resolution and computational efficiency. Smaller values of *k*, such as 2 or 3, yield low-resolution matrices that may fail to capture meaningful sequence patterns, while larger *k* values increase the dimensionality and sparsity of the matrix, leading to higher computational costs without a proportional gain in the biological signal [18]. A 6-mer provided a sufficiently informative context for modeling sequence features relevant to splice site prediction. The matrix was initialized to zero; for each *k*-mer occurrence, the corresponding cell count was increased by one. After scanning the sequence, the counts were normalized to [0,1], and a viridis color map [19] is applied. The final output is a 64 × 64 pixel RGB image.

For example, for the 6-mer ACGTAC (A = 00, C = 01, G = 10, T = 11, A = 00, C = 01), the position is:

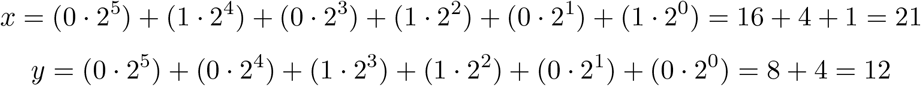

which maps ACGTAC to position (21, 12).

In the resulting image, brighter color intensities correspond to *k*-mers that occur more frequently within the DNA sequence.

Figure 1 illustrates the complete FCGR encoding pipeline, starting from the initial CGR walk (panel a), followed by the construction of the *k*-mer frequency matrix (panel b), and ending in the normalized 64 × 64 FCGR image used as input for deep learning models (panel c).

**Figure 1.**
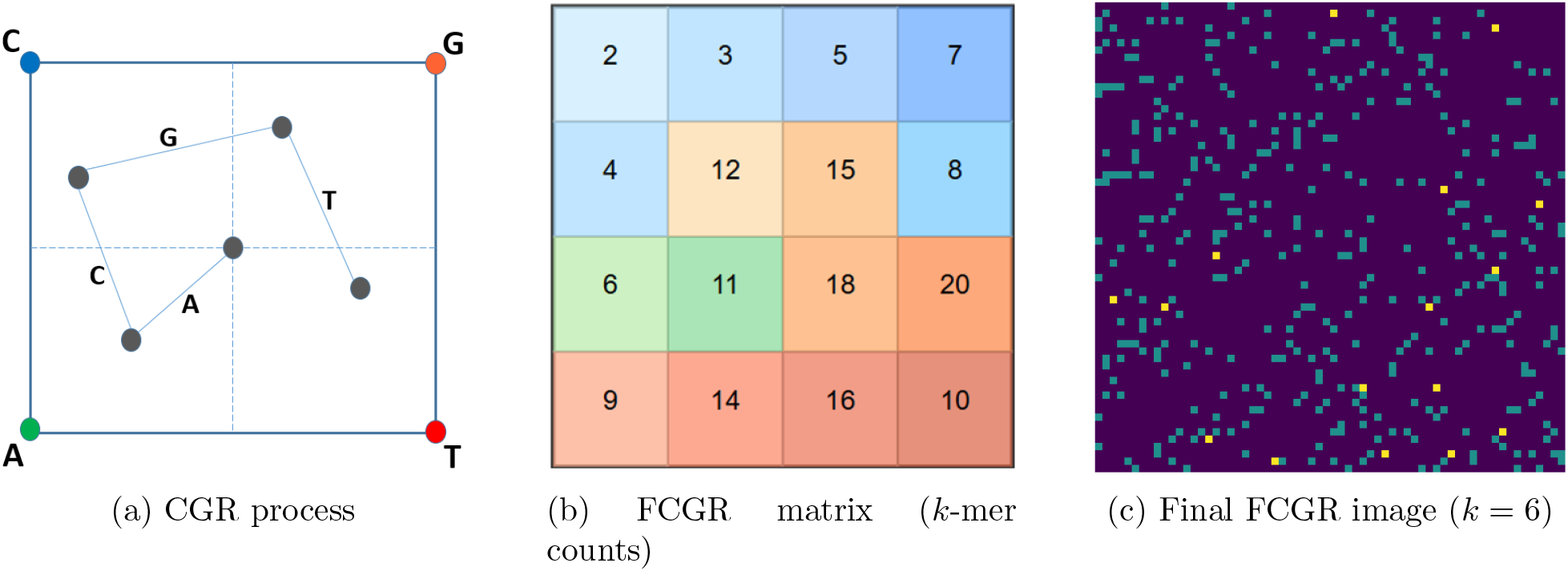
Overview of the Frequency Chaos Game Representation (FCGR) construction process. (a) CGR iteratively plots points in a square to reflect *k*-mer composition. (b) The resulting *k*-mer frequency mat rix is generated. (c) The matrix is then normalized and mapped to an RGB image.

### 2.3 Dinucleotide Fixed Color Pattern

We introduce a novel aproach, called DFCP, to encode DNA sequences into 2D images that preserve both local and long-range nucleotide relationships. Given a DNA sequence *S* = *s*_1_*s*_2_ *… s_n_*, DFCP constructs an *n* ×*n* matrix *M*, where each entry *M*_*i,j*_ represents the dinucleotide formed by (*s*_*i*_, *s*_*j*_):

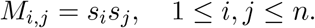

For example, for the short DNA sequence `ACGT`, the corresponding dinucleotide matrix is shown in Table 1.

**Table 1:** Example dinucleotide matrix for the sequence “ACGT”.

|  | A | C | G | T |
| --- | --- | --- | --- | --- |
| A | AA | AC | AG | AT |
| C | CA | CC | CG | CT |
| G | GA | GC | GG | GT |
| T | TA | TC | TG | TT |

Each unique dinucleotide (AA, AC, AG, …, TT) is then assigned a fixed RGB color according to a predefined mapping. The entire matrix is colored and rendered as an image.

In this study, sequences of length 402 nucleotides produce images with a resolution of 402 × 402. The diagonal entries represent local self-pairings, whereas the off-diagonal elements capture long-range nucleotide interactions. The resulting color pattern encodes the sequence context, enabling convolutional neural networks to detect complex spatial dependencies relevant to splice site prediction.

Figure 2 presents an example of a DFCP image alongside the fixed dinucleotide color legend used to encode pairwise nucleotide relationships between sequence positions.

**Figure 2.**
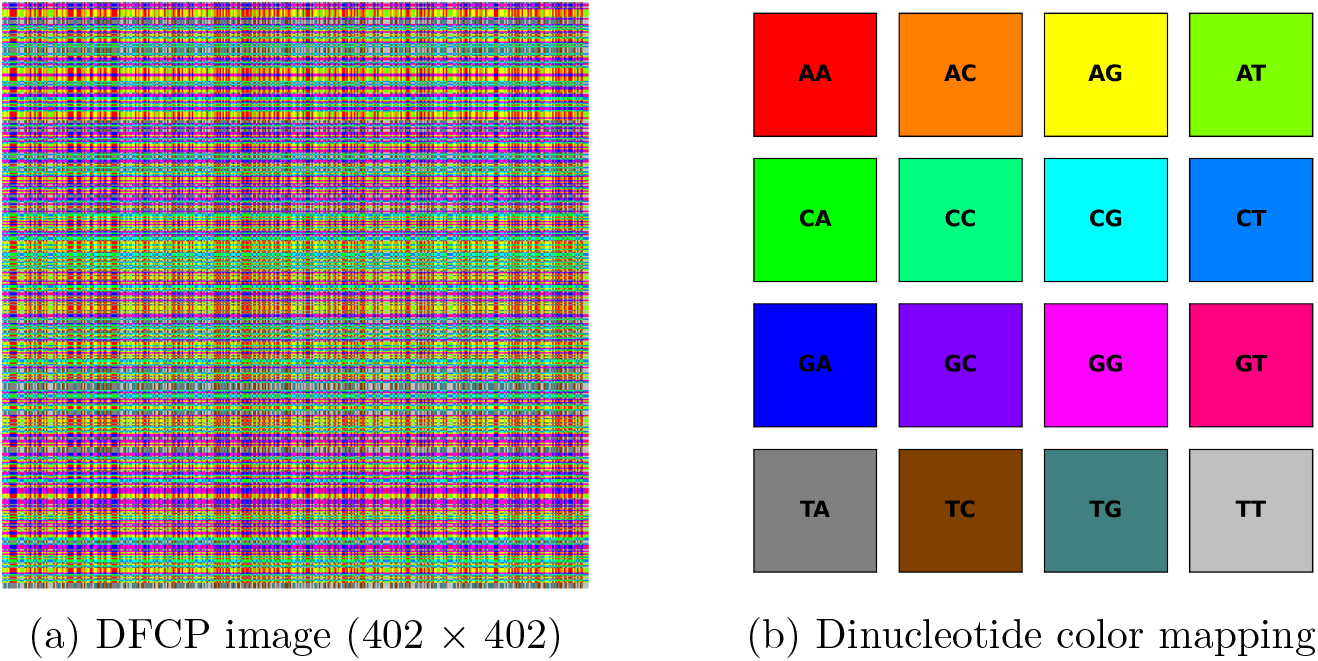
DFCP encoding. (a) A 402 × 402 image generated from positional dinucleotide pairings in a DNA sequence. (b) Fixed color mapping scheme used to represent each of the 16 possible dinucleotides.

### 2.4 Comparison of FCGR and DFCP

The two approaches offer complementary advantages for deep learning: FCGR emphasizes global motif patterns, while DFCP encodes fine-grained pairwise interactions. A direct comparison of their characteristics can be found in Table 2. This comparison highlights the complementary nature of the FCGR and DFCP representation methods, suggesting that integrating both representations may provide richer input features for deep learning models in splice site prediction.

**Table 2:** Comparison of the FCGR and DFCP encoding approaches.

| Characteristic | FCGR | DFCP |
| --- | --- | --- |
| Feature type | Frequency-based $k$-mer occurrence | Pairwise dinucleotide identity |
| Input encoding | Nucleotides mapped to binary codes; $k$-mer frequencies counted | Dinucleotides mapped to fixed RGB colors |
| Image size | $64 \times 64$ (for $k = 6$) | $402 \times 402$ (matching sequence length) |
| Information captured | Global $k$-mer composition and motif frequency patterns | Local and long-range nucleotide interaction patterns |
| Color meaning | Brighter intensity indicates higher $k$-mer frequency | Each dinucleotide is assigned a unique predefined color |
| Strengths | Captures overall sequence composition; compact image size | Preserves detailed pairwise relationships; retains spatial dependencies |

### 2.5 ResNet50 Model

We employed ResNet50 [16], a deep convolutional neural network widely used for image classification. Its residual connections enable effective training of deep architectures by mitigating vanishing gradients, making it well suited for learning complex spatial patterns in visual encodings of genomic data. Using binary cross-entropy loss and the Adam optimizer, we trained a ResNet50 model from scratch for 20 epochs, once on the FCGR dataset and once on the DFCP dataset. Training and evaluation were conducted separately for each species and splice site type to enable a systematic comparison of encoding strategies across biological contexts.

## 3 Results

### 3.1 Performance Evaluation

We evaluated the effectiveness of our models using four standard classification metrics: Accuracy, Precision, Recall, and F1-score. Let *TP, TN, FP*, and *FN* denote the number of true positives, true negatives, false positives, and false negatives, respectively. The metrics are defined as:

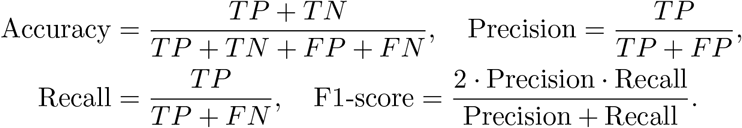

As shown in Table 3, DFCP consistently outperformed FCGR in all tasks and species. For *Arabidopsis thaliana*, DFCP achieved substantial improvements, particularly in donor site prediction, with an accuracy of 0.9294 compared to 0.7750 for FCGR and an F1-score of 0.9267 compared to 0.7841. We observed similar gains in effectiveness for acceptor sites, where DFCP reached an accuracy of 0.9186 and an F1-score of 0.9158, while FCGR achieved 0.7723 and 0.7826, respectively. In *Homo sapiens*, DFCP also led to superior results. For donor sites, the model reached an accuracy of 0.9573 and an F1-score of 0.9568, compared to 0.7767 and 0.7857 using FCGR. We also noted improvements for acceptor sites (accuracy: 0.9437 vs. 0.7484; F1-score: 0.9444 vs. 0.7529).

**Table 3:** Effectiveness of donor and acceptor splice site prediction in *Arabidopsis thaliana* and *Homo sapiens* for FCGR and DFCP image representations.

| Species | Encoding | Accuracy | Precision | Recall | F1-score |
| --- | --- | --- | --- | --- | --- |
| *Arabidopsis* Donor | FCGR | 0.775 | 0.7535 | 0.8174 | 0.7841 |
|  | DFCP | 0.9294 | 0.9627 | 0.8934 | 0.9267 |
| *Arabidopsis* Acceptor | FCGR | 0.7723 | 0.7485 | 0.82 | 0.7826 |
|  | DFCP | 0.9186 | 0.9474 | 0.8864 | 0.9158 |
| *Homo sapiens* Donor | FCGR | 0.7767 | 0.7552 | 0.8188 | 0.7857 |
|  | DFCP | 0.9573 | 0.9688 | 0.9450 | 0.9568 |
| *Homo sapiens* Acceptor | FCGR | 0.7484 | 0.7396 | 0.7668 | 0.7529 |
|  | DFCP | 0.9437 | 0.9332 | 0.9558 | 0.9444 |

Our results suggest that DFCP provides a more expressive encoding of both local motifs and long-range nucleotide interactions than FCGR, leading to more accurate and robust splice site classification across different species and splice site types.

### 3.2 Model Interpretability

Beyond the quantitative improvements observed with DFCP, we investigated the interpretability of the model predictions using saliency maps [20] and Grad-CAM [21]. Saliency maps highlight regions of the input that most influence model output by computing input gradients, while Grad-CAM identifies class-discriminative regions by weighting the activation maps of the final convolutional layer. The differences in interpretability between FCGR and DFCP can be explained by the structural properties of the two encoding schemes.

For FCGR, we mapped 6-mers to exact spatial positions in the 64 × 64 image using a deterministic coordinate function based on chaos game representation. Each nucleotide was represented by a 2-bit binary code (A = 00, C = 01, G = 10, T = 11), with the least and most significant bits determining the *x* and *y* coordinates, respectively. This enabled the precise location of important 6-mers on the Grad-CAM and the saliency heatmaps shown in Figure 3.

**Figure 3.**
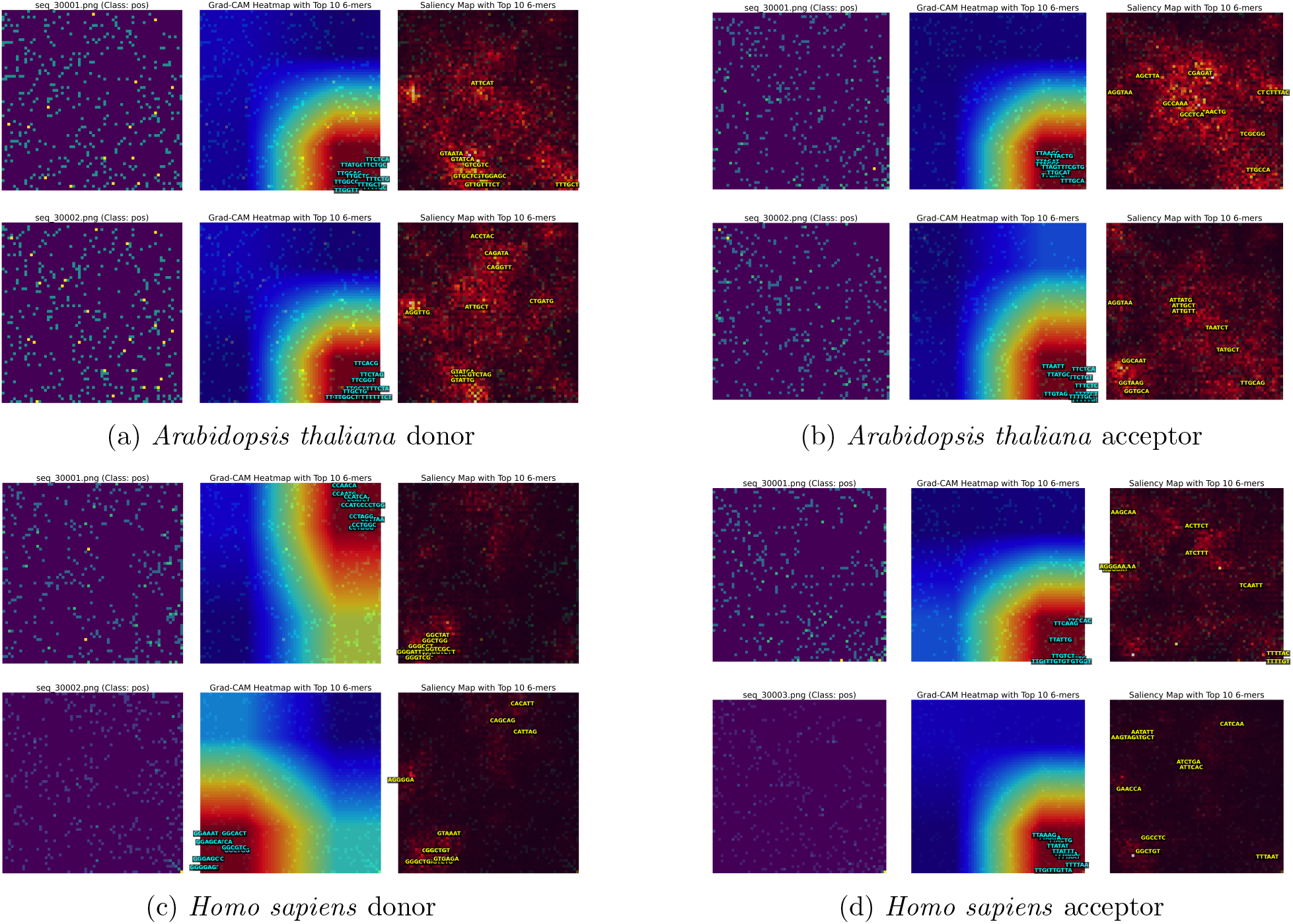
Interpretability results on FCGR image representations. For each task, we show the original input image, saliency map, and Grad-CAM heatmap for a correctly classified positive (true splice site) sample.

For *Arabidopsis thaliana* splice site prediction, both donor (Figure 3a) and acceptor (Figure 3b) tasks showed consistent recognition of TT-rich motifs. Grad-CAM emphasized polypyrimidine tract sequences including `TTGCAG`, `TTGGCC`, and `TTTGCA` for donors, and `TTATGT`, `TTGATC`, and `TTGCAT` for acceptors. Saliency maps highlighted diverse regulatory elements such as `CAGGTT`, `AGGTAA`, and `TATGCT` in donors, and `TTATAT`, `ACTTTG`, and `TTCTGT` in acceptors, reflecting sensitivity to both polypyrimidine tracts and auxiliary regulatory sequences characteristic of plant splice sites.

For *Homo sapiens* splice site prediction (Figure 3c and Figure 3d), the model demonstrated species-specific recognition patterns. In particular, Grad-CAM analysis of donor sites revealed activation of GC-rich motifs such as CCGGAC, GGACCT and `GGAGGG`, while saliency maps identified purine-rich sequences including AGGGTT, AGTAGG, and `AGGGGG`. Human acceptor sites showed mixed patterns, with Grad-CAM highlighting both TT-rich (TTGTCT, TTATGG) and GG-rich motifs (GGAAGG, GGGTTG), while saliency maps emphasized core acceptor signals like AGATGA alongside regulatory sequences such as `AGGAGT` and `TTTAGT`.

For DFCP, prominent horizontal activation bands were consistently observed near the center of the images across both donor and acceptor splice site tasks, as shown in Figure 4. These bands reflect the attention of a model on dinucleotides positioned around the splice junction. In donor sites, highlighted dinucleotides included `GT, GA, GC`, and `GG`, which are commonly associated with canonical donor motifs. In acceptor sites, the model frequently focused on `AG, CA, AA`, and `AC`, reflecting known sequence patterns near acceptor regions. These consistent spatial responses indicate that DFCP enables the model to learn biologically relevant motifs along with their surrounding sequence context. To better interpret these learned features, each 402 × 402 DFCP image was divided into 6 × 6 equal tiles of 67 × 67 pixels. Within each tile, Grad-CAM and saliency heatmaps were overlaid with the corresponding dinucleotide labels fixed at their spatial coordinates. Figure 5 provides an example derived from an *Arabidopsis thaliana* sequence containing a donor splice site.

**Figure 4.**
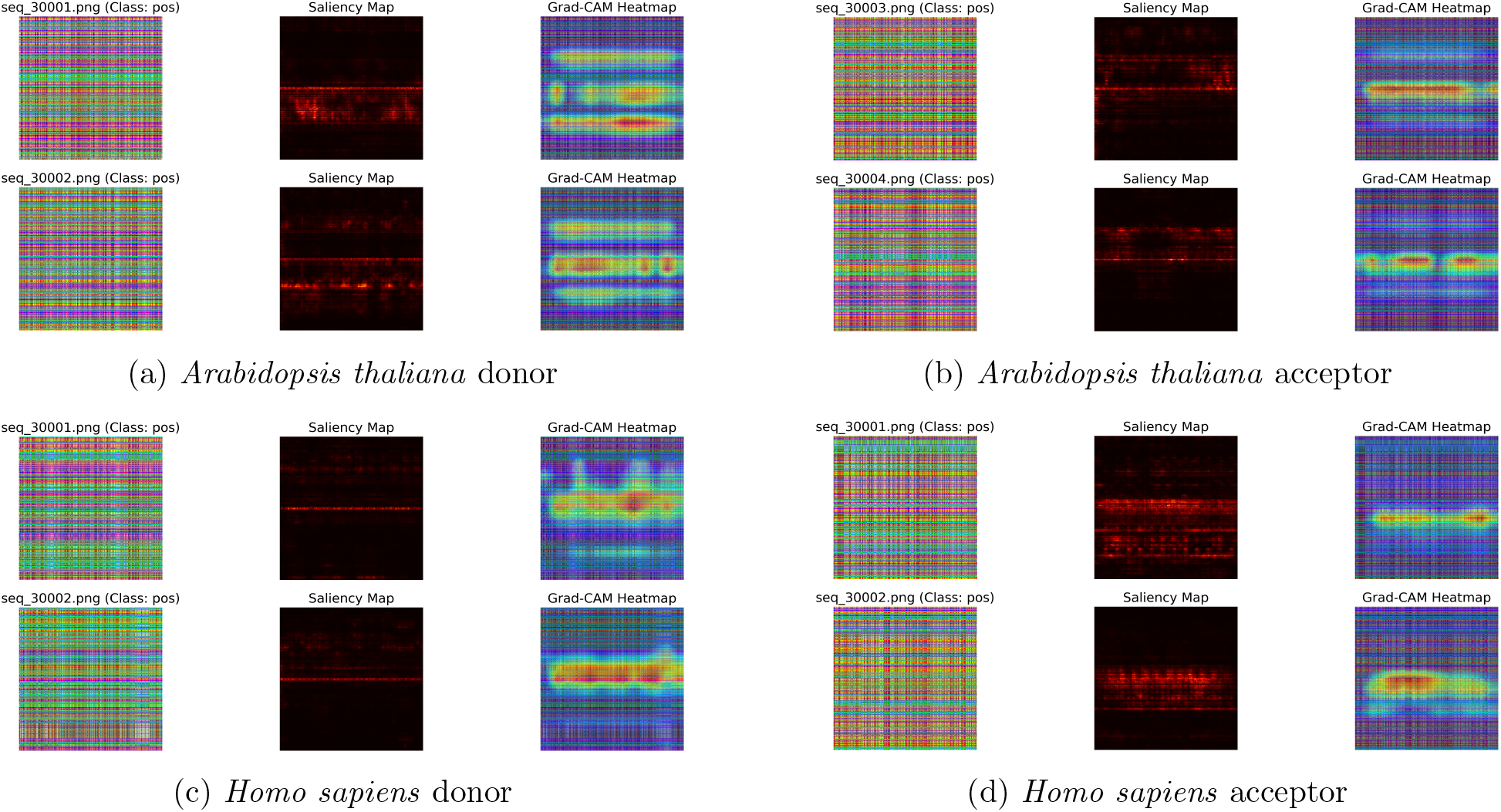
Interpretability results on DFCP image representations. For each task, we show the original input image, saliency map, and Grad-CAM heatmap for a correctly classified positive (true splice site) sample.

**Figure 5.**
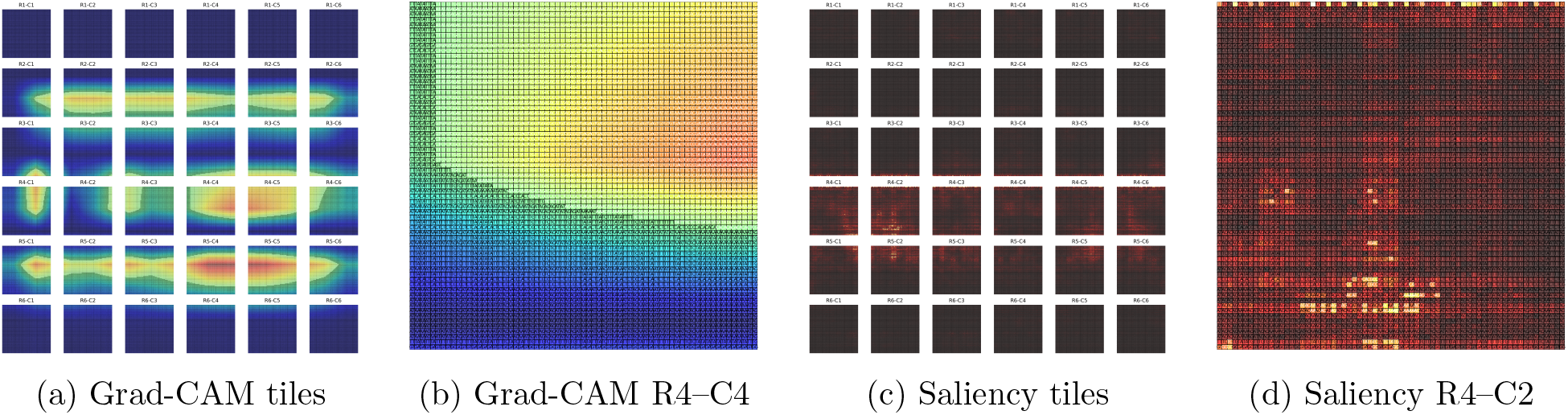
Grad-CAM and saliency visualizations for a DFCP input image derived from an *Ara-bidopsis thaliana* donor splice site sequence.

In Figure 5a, the Grad-CAM activation is shown across the full image with all 36 labeled tiles, while Figure 5b zooms into row 4, column 4, where strong model attention appears over key dinucleotide combinations near the splice junction. Similarly, Figure 5c presents the full saliency map, and Figure 5d offers a focused view on row 4, column 2, highlighting the dinucleotides contributing most to model predictions. These examples demonstrate that DFCP is effective in preserving and revealing spatially organized, interpretable signals critical to splice site recognition.

## 4 Conclusions and Future Work

We introduced DFCP, a novel visual encoding method for representing DNA sequences as images, and compared its effectiveness to the established FCGR method for splice site prediction using deep learning. By converting DNA sequences into image representations and applying a ResNet50 model, we demonstrated that DFCP consistently outperforms FCGR across donor and acceptor splice site classification tasks in both *Arabidopsis thaliana* and *Homo sapiens*. Interpretability analyses further revealed that DFCP yields more spatially coherent attention patterns, consistent with the expected positional organization of sequence features around splice sites. This behavior can be attributed to DFCP preserving both positional and compositional information in the encoded sequences.

In future work, we aim to extend our approach to a broader range of genomic tasks and species, and to explore its applicability with other neural network architectures. Additionally, we plan to further analyze saliency and Grad-CAM maps to uncover conserved motifs and dependencies, improving the interpretability and reliability of deep learning models in genomics.

## Data Availability

The dataset (DNA sequences) used in this study are available at https://github.com/XueyanLiu-creator/DRANetSplicer/tree/main/data/dna_sequences and the code at https://github.com/EspoirKabanga/SpliceImage.

